# A Murine Database of Structural Variants Enables the Genetic Architecture of a Spontaneous Murine Lymphoma to be Characterized

**DOI:** 10.1101/2025.01.09.632219

**Authors:** Wenlong Ren, Zhuoqing Fang, Egor Dolzhenko, Christopher T. Saunders, Zhuanfen Cheng, Victoria Popic, Gary Peltz

## Abstract

A more complete map of the pattern of genetic variation among inbred mouse strains is essential for characterizing the genetic architecture of the many available mouse genetic models of important biomedical traits. Although structural variants (SVs) are a major component of genetic variation, they have not been adequately characterized among inbred strains due to methodological limitations. To address this, we generated high-quality long-read sequencing data for 40 inbred strains; and designed a pipeline to optimally identify and validate different types of SVs. This generated a database for 40 inbred strains with 573,191SVs, which included 10,815 duplications and 2,115 inversions, which also has 70 million SNPs and 7.5 million insertions/deletions. Analysis of this SV database identified an SV that is one component of a bi-genic model for lymphoma susceptibility in SJL mice, which provides mechanistic insight into the genetic basis for susceptibility to murine (and potentially human) lymphomas.

## INTRODUCTION

The laboratory mouse has been the premier model organism for biomedical research, and the large number of phenotypically well-characterized inbred strains has enabled genetic factors for important biomedical traits to be identified using murine genetic models ^1–3^. However, mouse genetic discovery is critically dependent upon having a complete map of genetic variation among these strains. While SNPs and small indels have been extensively characterized in mouse strains ^4–6^, the lack of a comprehensive database of structural variation has limited our ability to fully analyze and interpret the mouse genome. Prior efforts to characterize murine structural variants (SVs) (i.e., genomic alterations >50 bp in size) have included only a few strains ^7^ and relied on short-read sequencing (SRS) ^8,9^, which has a limited ability to detect SVs in repetitive regions of the genome. Long-read sequencing (LRS) platforms, which produce reads of length ≥20kb, have improved our ability to identify SVs, especially in repeat-rich genomic regions^10–12^. LRS has doubled the estimated number of SVs in the human genome versus prior SRS estimates ^10^.

Our prior analysis of six strains with LRS revealed that: (i) SVs are very abundant (4.8 per gene), which indicates that they are likely to impact genetic traits; and (ii) as in human studies, the SVs previously identified using SRS^13^ accounted for only 25% of those identified by our LRS analysis ^14^. A recent analysis of SVs in 14 inbred strains used LRS ^15^; but this analysis primarily reported only deletions and insertions. Sequencing and alignment artifacts, along with a heavy reliance on heuristics limit the ability of many existing programs to accurately identify additional types of SVs. We found that characterization of duplications or inversions was particularly problematic, even when murine LRS was analyzed ^14^. To comprehensively characterize SVs across the mouse genome, we sequenced 40 inbred mouse strains using high-accuracy PacBio HiFi long reads. Simulations were used to evaluate the performance of several SV detection methods for identifying different types and sizes of murine SVs. In addition to state-of-the-art alignment ^16,17,18^ and assembly-based ^19^ heuristic methods, we also evaluated a recently developed deep-learning method (Cue ^20^) for SV detection. Based upon these simulation results, we designed a custom pipeline to characterize a broader set of SVs from these 40 strains, which included deletions (DEL), insertions (INS), duplications (DUP), and inversions (INV) of varying size. This approach generated a comprehensive database of 573,191SVs among 40 inbred strains that includes a significant number of novel DUPs, INVs and large INS.

The utility of this SV database was demonstrated by identifying a genetic susceptibility factor for an unusual lymphoma that spontaneously appears in SJL mice ^21–23^. These tumors originate in B cell germinal centers and are of interest because some of their features resemble those seen in one type of human non-Hodgkins lymphoma ^24^. SJL lymphomas contain activated T cells, which produce the cytokines required for lymphoma propagation *in vitro* ^25^. One susceptibility factor was identified as an endogenous retrovirus (***Mtv29***) in the SJL genome that encodes a tumor associated antigen (**vSAg29**) ^26,27^, which stimulates a subset of CD4 T cells ^28^ to produce the cytokines required for lymphoma development ^29^. However, a second genetic susceptibility factor also must contribute because: (i) a strain (MA/My) that expresses vSAg29 does not develop (or has a very low incidence of) lymphoma ^30^; and (ii) analyses of SJL intercross progeny indicated that one autosomal dominant genetic factor is present in multiple other strains, which suppresses lymphoma development ^31,32^. Although this tumor suppressor had not been identified in the 55 years since this lymphoma was described in 1969, our AI mouse genetic discovery pipeline ^33^ identified this second genetic factor as an SJL-unique SV that ablates a tumor suppressor.

## RESULTS

### Genomic sequencing and SNP/INDEL identification

Genomic sequencing was performed using a PacBio Revio instrument with the HiFi system, which achieves a median read accuracy reaching 99.9% ^34^, to generate LRS from 40 inbred strains. A total of 3.54 TB of sequence was generated, with an average of 88.5 GB (30x coverage) per strain (**Table S1**). Since they are commonly used in genetic models, we separately report on SNPs, Indels and SVs present the 35 classic inbred strains and those present in all 39 sequenced strains, which includes four wild derived strains (CAST, SPRET, MOLF, WSB). Using DeepVariant (v1.6.1) ^35^, 70,051,144 SNP sites and 7,540,144 sites with insertions or deletions (INDELs) were identified in the 39 strains (vs the C57BL/6 reference genome GRCm39). Consistent with our previous finding that the four wild-derived strains had patterns of genetic variation that were distinct from the 35 commonly used classical inbred strains ^36^, most of the SNPs (64.7M or 92%) and INDELs (6.8 M or 91%) were present in the 4 wild-derived strains, which also had most of the SNP or INDEL alleles that were present in only one strain. There were 21,331,225 SNP and 2,290,861 INDEL sites in the 35 classical inbred strains; the alleles in 5,286,543 SNPs and 696,031 INDELs were only present in the classic inbred strains; and 4.7M SNPs and 0.52M INDELs had alleles that were present in only a single classic inbred strain (**Figure S1**). The missing allele call rates for SNPs (0.005) and INDELs (0.03) were very low. To ensure that the strains were correctly labelled, we examined the output of our AI mouse genetic analysis pipeline ^33^ using the new SNP and INDEL databases. This AI pipeline, which uses novel machine learning methodology, was developed to rapidly analyze murine genetic model data, and identify high probability causal genetic factors for an analyzed trait. This pipeline first identifies candidate genes by genetic association analysis and then uses a novel graph neural network-based method to assess the 29M published papers in the biomedical literature to analyze candidate gene-phenotype relationships. The previously identified causative genetic factors for four traits ^14,33,36,37^ (two of which were caused by SNP alleles and two were caused by INDELs) were identified by the AI using the new SNP and INDEL databases (**Figures S2 and S3**). Thus, the strains used to generate the SNP and INDEL alleles in the new database are correctly labeled. We compare the SNPs identified here using the PacBio HiFi LRS with the SNPs previously identified using short-read sequencing (SRS) ^5^. Among the SNPs in the 39 strains present in both datasets, 85% of the SNPs overlapped (**Figure S4**). LRS detected 10% more SNPs than the prior study, which is consistent with the improved performance associated with the increased sequencing depth and the use of HiFi LRS methodology in this study. Moreover, the IGV for 200 randomly selected SNPs, which were among 9M that were only identified by the LRS methodology, were examined. Only 4 of these 200 SNPs (2%) had minor issues that involved misclassification of genotypes (homozygous vs heterozygous) or of complex allelic patterns; and all of these were in SNPs present in wild-derived strains (Figure S4). Since the 69.9M SNPs that were jointly identified by analyses using different sequencing methods can be considered as validated, and the LRS-only SNPs have a 98% true positive rate; this SNP database has an overall true positive rate of 99.7%.

### Assessing SV identification programs

Since SV analysis programs vary in their ability to identify different types of SVs ^38,39^, we performed an extensive set of simulations to examine the ability of five programs (Cue ^20^, Sawfish (v0.10.0) ^18^, Sniffles2 (v2.3.2) ^16^, PBSV (v2.9.0) ^17^, and Dipcall (v0.3) ^19^) to detect SVs that were artificially inserted into the mouse genomic sequence. We separately assessed their ability to identify small (<1 kb) and large (1-100 kb) DELs, INSs, INVs and DUPs (**Table 1**). The simulation results were used to design the SV pipeline used to assemble this SV database (**Figure 1**). Whenever possible, we used a consensus-based SV identification strategy. However, when only a single caller achieved high recall in the simulations (i.e., large INSs), or when low agreement was observed among the programs when the actual data was analyzed (i.e., INVs) (**Figures 2A and S5A**), the consensus strategy was replaced with one where the individual predictions obtained from one or more high recall tools were selected if they were validated by another program (VaPoR ^40^) or by visual inspection. The evaluation metrics and parameter thresholds (especially breakpoint stringency and sequence similarity) can influence the results obtained from SV detection tools ^38^. Therefore, we performed simulations using three different parameter settings (default, stringent, relaxed) and evaluated SV caller performance for identification of different types and sizes of SVs (**Tables S2 and S3**). We found that the SV identification programs we selected were robust to variation in breakpoint matching criteria since only minor differences were observed when three different parameter configurations were used. Overall, our results indicate that DEL and INS can be reliably identified, but improved methods for identification and validation of DUPs and INVs are needed.

**Figure 1.**
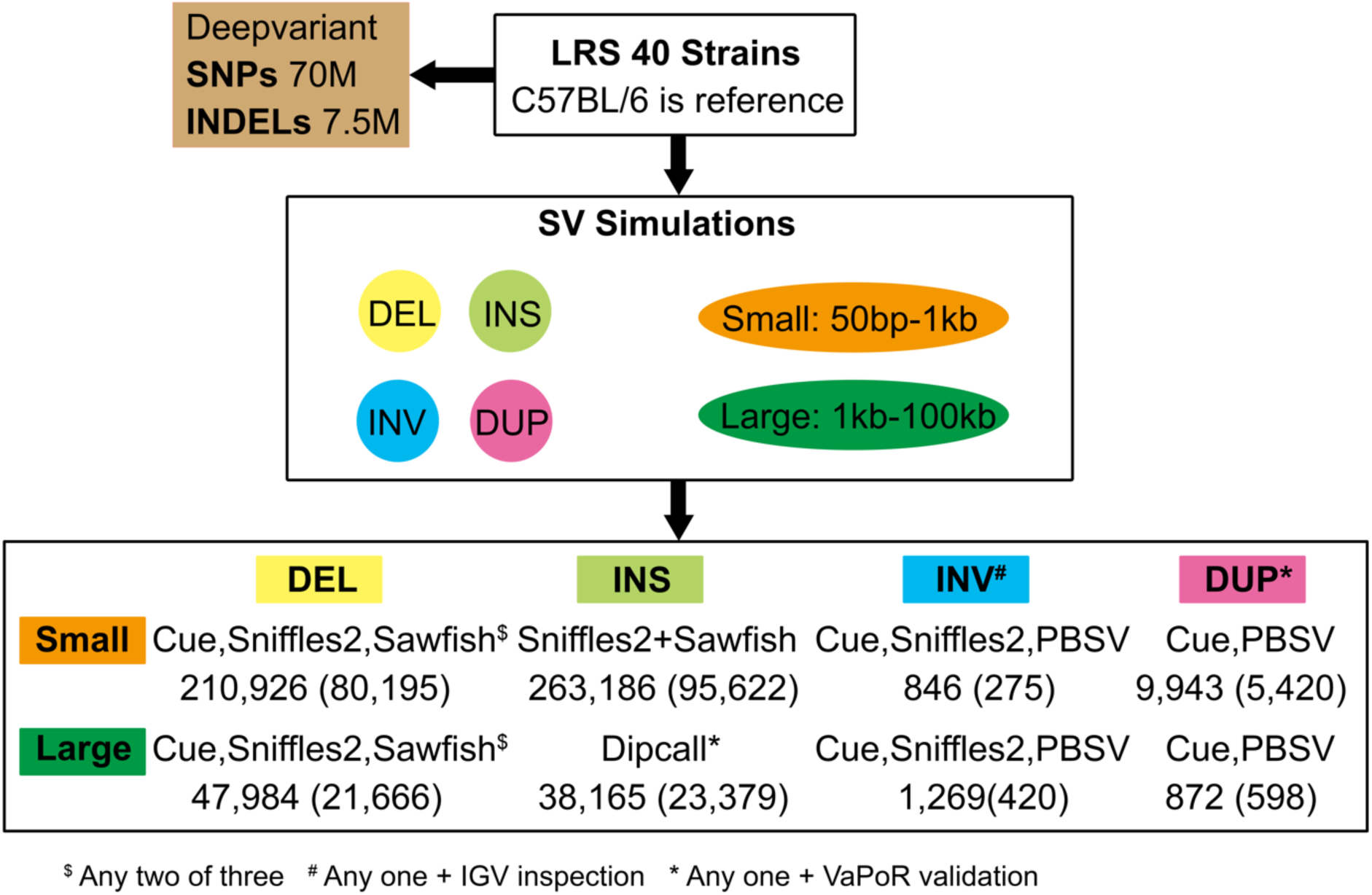
Summary of the pipeline used to analyze the genomic sequences of 40 inbred mouse strains to produce the SNP, INDEL, and SV databases and their content. Long Range Sequencing of 40 inbred strains was performed, and the C57BL/6 sequence was used as the reference sequence. Deepvariant was used to identify alleles for 70M SNPs and 7.5 million insertions- deletions in the 40 strains. SVs were separately analyzed based upon their size (small, 50 to 1000 bp; large, 1 to 100 KB) and type (DEL, INS, INV, DUP). The simulation results (Table 1) and the overlap among SVs identified by the programs were analyzed to produce the SV identification procedures shown in this figure. There were 210,926 small DELs, 47984 large DELs, and 263,186 small INS that were jointly identified by two of the three analysis programs (Cue, Sniffles2, Sawfish). Since none of these programs could reliably identify large INS, the genomic sequences were individually assembled and Dipcall was used to identify 38,165 large INS that were verified using the VaPoR program. To identify small and large INVs, any INV identified by Cue, Sniffles2, or PBSV that was manually verified using the IGV program was reported. The 9,943 small and 872 large DUPs identified by Cue or PBSV were validated by the VaPoR program. The numbers within parenthesis indicate the number of each type of SV present in the 35 commonly used inbred strains.

**Figure 2.**
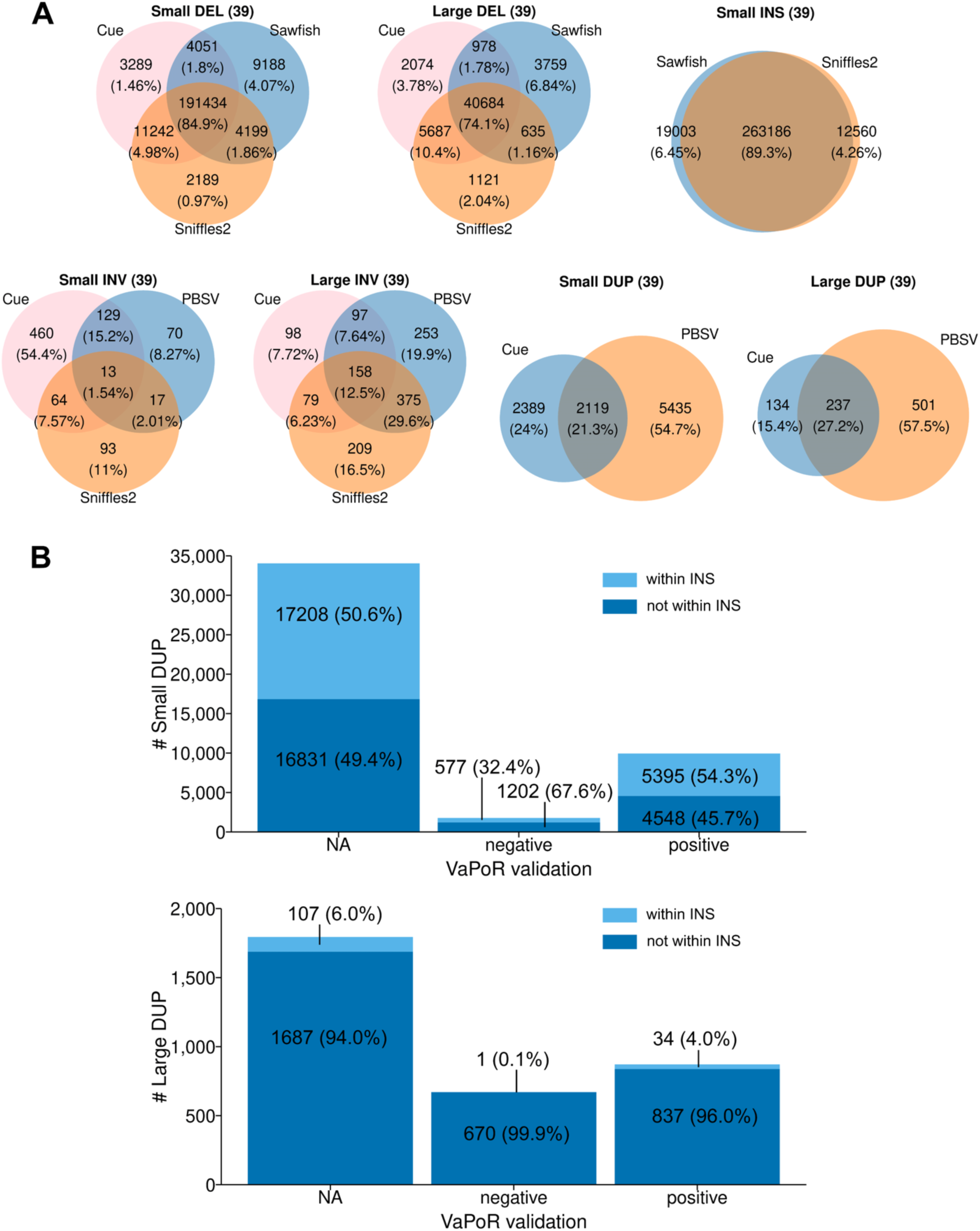
Assessing the performance of different SV identification programs. (A) SV identification programs have an acceptable level of agreement for identification of small (50 – 1000 bp) and large (1 kb – 500 kb) deletions (DELs), and for small insertions (INSs). The results obtained from analysis of the genomic sequences all 39 inbred strains by each indicated program are shown in separate Venn diagrams. The number of SVs identified by an individual (or a combination of) programs are shown within the circles and the percentages are shown in parenthesis. In contrast, there was substantial divergence in the INVs identified by Cue, PBSV and Sniffles2. Because of this, the INVs identified by each program were verified by manual inspection using the IGV program. (B) The Vapor validation results for DUPs identified by Cue or PBSV. Most of the DUPs identified by Cue or PBSV could not be assessed by VaPoR (represented by NA). The DUPs that passed or failed the VaPoR validation are represented as positive or negative, respectively. We also examined whether DUPs were found within an identified INS: 54% of validated small DUPs were within an INS, but only 4.0% of validated large DUPs were within an INS.

**Table 1.** The results of simulations assessing the ability of Cue, Sniffles2, Sawfish, PBSV and Dipcall to identify various types of SVs are shown. For each simulation, 1000 SVs of a single type (DEL, INS, INV, DUP) with a restricted size range [small (50 - 1000 bp) or large (1 - 100 KB)] were artificially inserted at random positions into the GRCm39 reference sequence. The precision (P), recall (R) and F1 statistic were calculated based upon the ability of each program to detect each type of inserted SV. Cue, Sniffles2, Sawfish and PBSV analyzed Hifi LRS (30x coverage) that were aligned with minimap2. The simulations were performed using PBSIM3. The genomic sequences analyzed by Dipcall were assembled using HiFiasm. NA: not assessed. W* and W^$^: withdrawn.

|  |  | Cue | Sniffles2 | Sawfish | PBSV | Hifi-asm<br>Dipcall |
| --- | --- | --- | --- | --- | --- | --- |
|  |  | *LRS*<br>*alignment+DL* | *LRS*<br>*alignment* | *LRS*<br>*alignment* | *LRS*<br>*alignment* | *LRS*<br>*assembly* |
| **DEL**<br>**(50-1000bp)** | **P** | 1 | 1 | 1 | 1 | 0.99 |
|  | **R** | 0.994 | 0.999 | 1 | 1 | 0.702 |
|  | **F1** | 0.996 | 0.9995 | 1 | 1 | 0.821 |
| **DEL**<br>**(1-100kb)** | **P** | 1 | 0.991 | 0.989 | 0.997 | 1 |
|  | **R** | 0.998 | 0.548 | 0.995 | 0.648 | 0.494 |
|  | **F1** | 0.999 | 0.706 | 0.992 | 0.785 | 0.661 |
| **INS**<br>**(50-1000bp)** | **P** | NA | 0.998 | 1 | 1 | 0.987 |
|  | **R** | NA | 0.978 | 0.982 | 0.979 | 0.811 |
|  | **F1** | NA | 0.988 | 0.991 | 0.989 | 0.89 |
| **INS**<br>**(1-100kb)** | **P** | NA | 1 | 1 | 1 | 1 |
|  | **R** | NA | 0.182 | 0.358 | 0.13 | 0.936 |
|  | **F1** | NA | 0.308 | 0.527 | 0.23 | 0.967 |
| **INV**<br>**(50-1000bp)** | **P** | 0.999 | 0.908 | W* | 0.967 | NA |
|  | **R** | 0.909 | 0.434 | W* | 0.758 | NA |
|  | **F1** | 0.952 | 0.587 | W* | 0.85 | NA |
| **INV**<br>**(1-100kb)** | **P** | 0.999 | 0.994 | W* | 0.994 | NA |
|  | **R** | 0.999 | 0.998 | W* | 0.901 | NA |
|  | **F1** | 0.999 | 0.996 | W* | 0.9454 | NA |
| **DUP**<br>**(50-1000bp)** | **P** | 0.997 | 0 | W\$ | 0.792 | NA |
| | **R** | 0.976 | 0 | W\$ | 0.795 | NA |
| | **F1** | 0.987 | 0 | W\$ | 0.793 | NA |
| **DUP**<br>**(1-100kb)** | **P** | 0.999 | 0.92 | 0.956 | 0.986 | NA |
|  | **R** | 0.993 | 0.916 | 0.762 | 0.982 | NA |
|  | **F1** | 0.996 | 0.918 | 0.848 | 0.984 | NA |

Nevertheless, an increased number of SVs, which includes DUPs and INVs, were identified by this pipeline.

### Characterization of SVs

A total of 210,926 small (50 to 1000 bp) DEL SVs were identified in the 39 inbred strains (vs the C57BL/6 reference), and 80,195 of them were present in the 35 classical inbred strains (Figure 1). There were 47,984 and 21,666 large (1 to 100 KB) DELs identified in the 39 or 35 inbred strains, respectively (Figures 2A and S5A). Like the SNP alleles, most small (61%) and many large (53%) DELs are only present in the four wild-derived strains. There were 263,186 and 95,622 small INS SVs in the 39 or 35 inbred strains, respectively (Figures 2A and S5A). Dipcall identified 38,165 or 23,379 VaPoR-validated large INS in all 39 strains or in the 35 classical inbred strains, respectively. Consistent with the simulation results, 94% (38,165 out of 40,460) of the large INS identified by Dipcall were validated by VaPoR, and most large INS that could not be validated had sequence complexity that precluded VaPoR analysis (i.e., rated as not assessable). The 846 (or 275) small INVs and 1,269 (or 420) large INVs were validated by visual inspection using the integrated genome viewer (IGV). In addition, the 9,943 (or 5,420) small DUP and 872 (or 598) large DUP in all 39 (or 35 classical) inbred strains were also validated using VaPoR. Most of the small and large DUPs identified by the analysis programs could not be assessed by VaPoR (**Figures 2B and S5B**). Among the 573,191SVs identified, the number of INS (n=301,351) and DEL (n=258,910) was far greater than DUPs (n=10,815) or INVs (2,115) (**Figure 3A**) in the 39 strains analyzed. SPRET, CAST/Ei and MOLF had the largest total number of SVs and of strain-unique SVs (**Figures 3B and 3C**). Three features of this analysis were notable. (i) Most small DUPs (54%) were within INS, while only 4.0% of the large DUPs were within INS (Figures 2B and S5B). (ii) Only a minority of the DUPs identified by Cue or PBSV could be validated by VaPoR (Figures 2B and S5B); visual inspection confirmed that sequence complexity made it difficult to assess these SVs. (iii) Most SV alleles were shared by three or fewer inbred strains (**Figures 3D and S6**), and were <1 kb in size (**Figure S7**).

**Figure 3.**
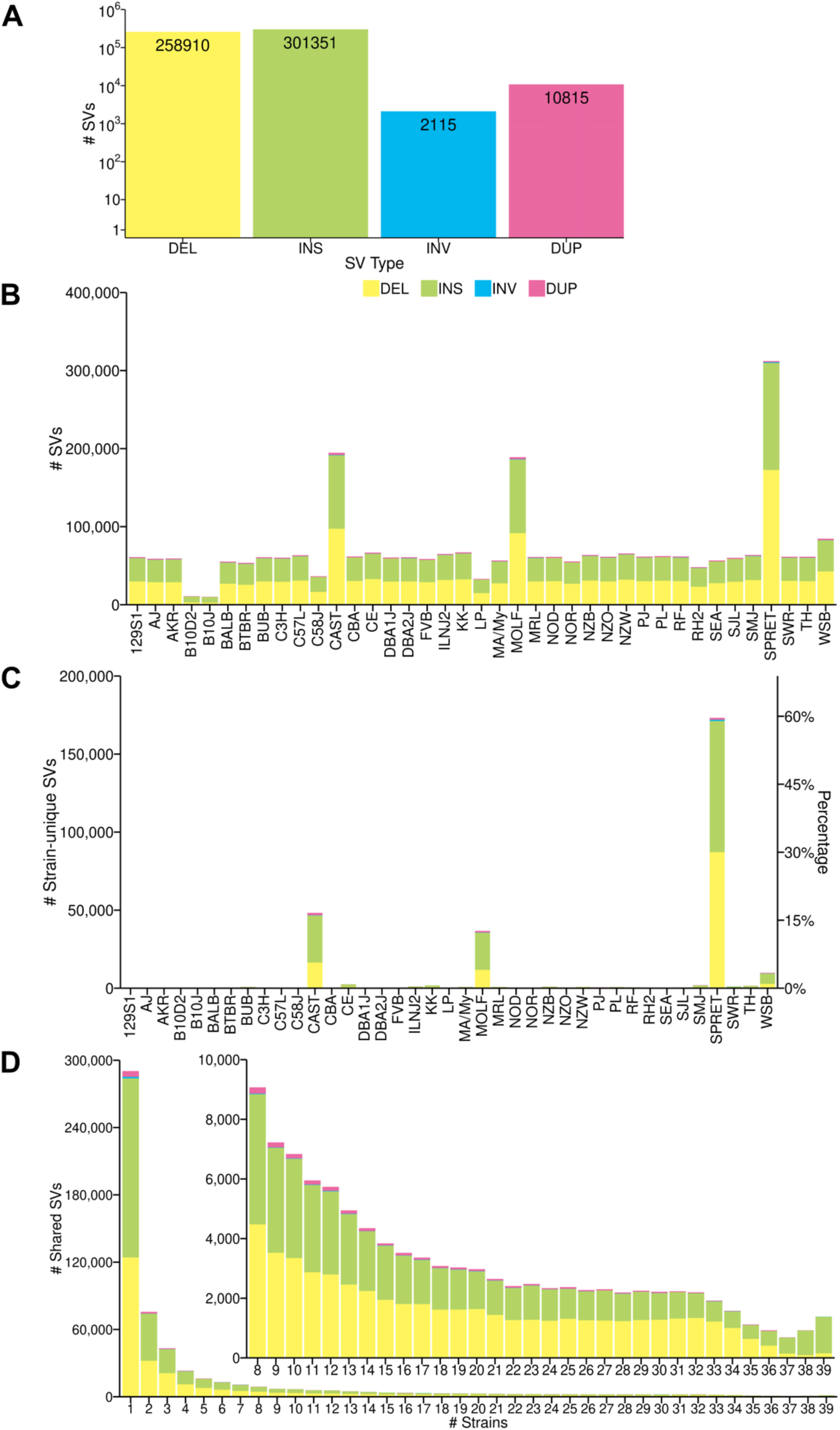
Characteristics or the SV alleles among the inbred strains. (A) The total number deletion (DEL), insertion (INS), inversion (INV) or duplication (DUP) SVs detected in all 39 strains are shown. (B) The number of each type of SV detected in each of the 39 strains are shown. Each type of SV is indicated by the color shown in A. (C) The number (left y-axis) and percentage (right y-axis) of strain-unique SVs detected in each of the 39 inbred strains is shown. Three wild derived strains (CAST, SPRET, MOLF) have most of the strain-unique SVs. (D) The number of strains with a shared SV allele are shown for all 39 inbred strains. Each type of SV is indicated by the color shown in A. Most of the minor SV alleles are shared by 1-3 strains. The inset graph shows the number of SV alleles shared by 8 or more strains.

We also compared the SVs identified by our analysis of 39 inbred strains with those identified in a prior study that analyzed SVs in 14 inbred strains ^15^ (**Figure 4, Table S4**). We identified 90,246 more DELs (53% increase) and 55,544 more INS (22% increase) than the prior study; and only 43% of the DELs and 35% of the INS were jointly identified by both studies. Given the increased number of strains that we analyzed, the increased number of DEL and INS identified by this study was expected. The non-overlapping DEL and INS SVs result from the two wild-derived strains (PWD/PhJ and PWK/PhJ) analyzed in ^15^, which were not among the four wild derived strains analyzed here. However, our study identified 10- and 25-fold more INVs and DUPs, respectively, than the prior study; and there was very little overlap in the INVs (2.4%) and DUPs (0.6%) identified by both studies. Our use of computational programs, which were selected based upon their performance in simulation studies for identifying INVs and DUPs, accounts for the increased number of INVs and DUPs identified by this study.

**Figure 4.**
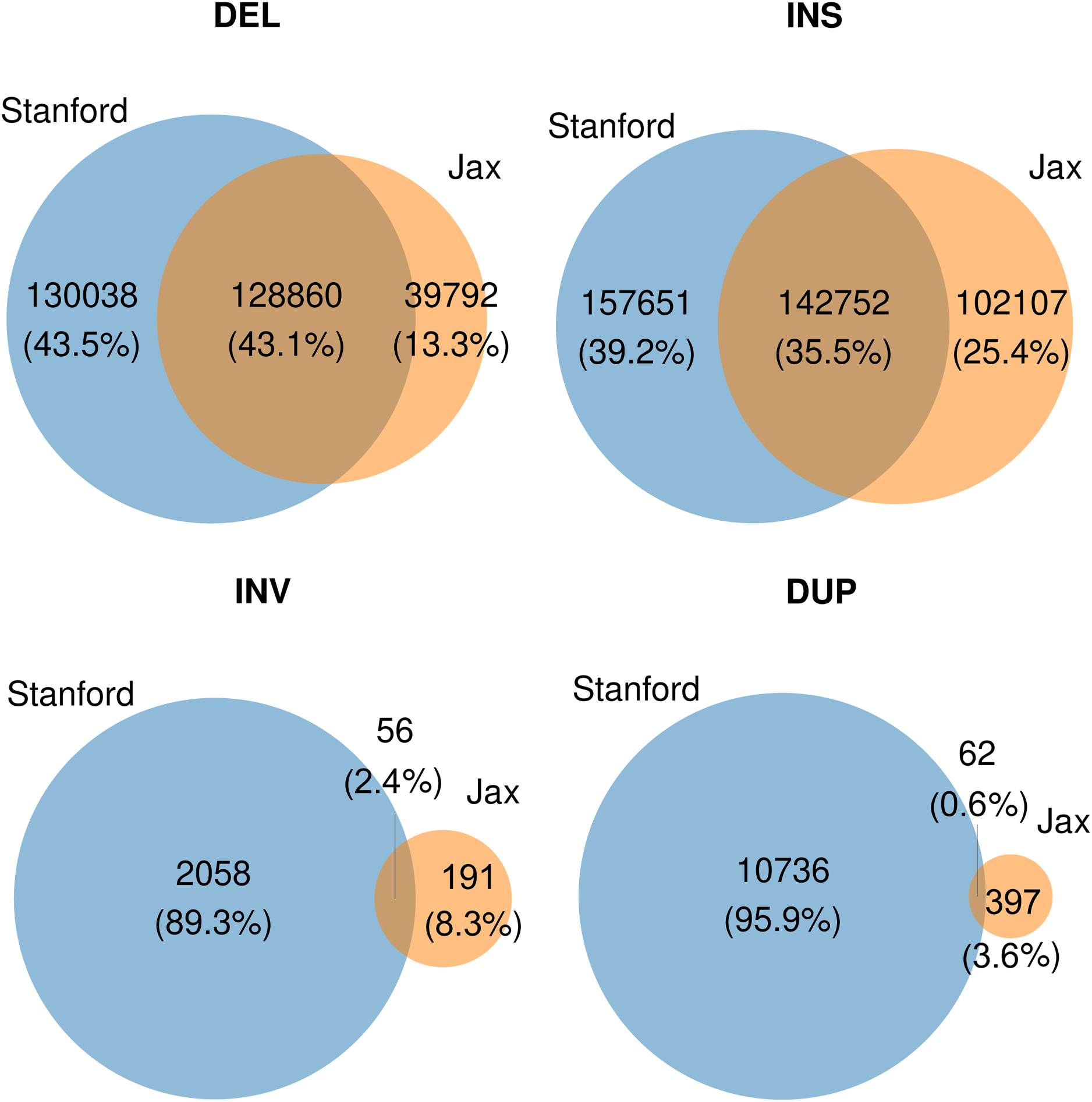
Venn diagrams comparing the number of SVs identified in this study (Stanford) from analysis of 39 inbred strains with those identified by the Jackson Laboratory’s (JAX) analysis of 14 inbred strains. The SV comparisons are separately categorized based upon whether they are a deletion (DEL), insertion (INS), inversion (INV), or duplication (DUP). The total number of each type of SV and the percentages are shown in parenthesis.

The variant effect predictor (VEP) program ^41^ was used to analyze SV impact (**Table S5**). Manual inspection of the high impact (protein coding) small INS (performed using the integrated genome viewer, IGV) revealed that 99.8% were true positives (2 false positives out of 801 analyzed), and 99.3% of the small DELs were true positives (5 false positive out of 705). Also, >99% of the large INS (1 false positive out of 518) and 98.6% of large DELs (4 false positives out of 293) were true positives (**Table S6**). To facilitate genetic discovery the genes with high impact SVs present in the 35 classical inbred strains are provided for small (n=705 in 654 unique genes) and large (n=293 in 270 unique genes) DELs, for small (n=801 SVs in 765 unique genes) and large (n=518 in 487 unique genes) INS, for all INVs, and for high impact DUPs in **Tables S7, S8, S9 and S10.** In summary, the 2,305 high impact SVs with alleles in the 35 classical inbred strains provides a set of highly curated genetic variants that could impact a substantial number of biomedical traits.

Although we have only a limited ability to interpret the impact of SVs located within intergenic and noncoding regions, chromatin is compartmentalized into topologically associating domains (**TADs**), which are megabase-sized genomic segments that are separated by boundary regions ^42,43^. TADs provide a regulatory scaffold for gene expression; they are linked with variation in gene expression because their structure facilitates enhancer-promoter interactions; and they insulate regions from the effect of other regulators ^44^. TADs are created by the binding of a DNA sequence-specific transcription factor (CCCTC binding factor or **CTCF**) to its consensus binding element. A multi-subunit protein (cohesin) then binds to CTCF to form the 3D loop-like structures that alter gene transcription within a domain. Of note, we found 1,877 DELs that contain a CTCF recognition sequence (CCGCGNGGNGGCAG) among the 35 inbred strains (**Table S11**). To determine if these DELs could impact chromatin structure, CTCF ChIP-Seq data from the ENCODE project ^45^ was examined to identify which of these CTCF recognition sequences were bound by CTCF. Among the 42 CTCF ChIP-Seq datasets that used mouse tissues or cell lines, CTCF binding occurred at 712 of the CTCF recognition sequences that were contained within the 488 deletion alleles. Since CTCF binding is critical for TAD formation, it is likely these deletions affecting these 712 CTCF recognition sequences could significantly alter chromatin structure; and by this mechanism could affect gene expression patterns and genetic traits.

### C57BL/6 sequence comparison

We report C57BL/6 alleles for each variant (based upon the GRCm39 reference sequence), which were identified by comparison with the other 39 strains. However, we also conducted a dedicated variant calling analysis, which compared our C57BL/6J data to the GRCm39 reference sequence for C57BL/6 produced by the Genome Reference Consortium. Even when a reference-based approach is used, subtle differences can be identified when sequences from the same strain are compared. We found very subtle increases in the number of SNPs (11.7K/70M or 0.017% of the total), INDELs (9.8K/7.5M or 0.13%), and in several types of SVs that were identified using our LRS data vs GRCm39. There was a 1.4% increase in the number of insertions (>90% small INS) identified using our LRS data vs the GRCm39 (**Table S12**). We suspect that the increased number of insertions identified using our LRS data is due to improved sequencing technology (HiFi LRS vs the short read sequencing that was used to compile GRCm39).

However, we can’t definitively determine whether sequencing technology differences or from genetic divergence between the C57BL/6 strains used by us and the Genome Reference Consortium. Since the GRCm39 is universally used by scientists, the sequence comparisons used to produce this database were performed relative to GRCm39.

### Identification of the 2nd lymphoma susceptibility factor

Based upon the hypothesis that SJL mice lack a tumor suppressor, the AI mouse genetic pipeline ^33^ was used to identify lymphoma-associated (MeSH Term: D008223) genes with high impact SV alleles uniquely present in SJL mice (i.e., absent in the 34 other classic inbred strains). The AI identified a 1641 bp SJL- unique deletion in *high mobility group A1b* (*Hmga1b),* which ablated the exon encoding the entire Hmga1b protein, as the candidate genetic factor (**Figure 5).** Hmga1b was the only gene with an SJL- specific high impact DEL that was directly associated with lymphoma. HMGA family members are low molecular weight proteins that bind to AT-rich regions in nuclear chromatin where they exert positive or negative effects on gene expression by enabling other transcription factors to bind at nearby sites ^46,47^.

**Figure 5.**
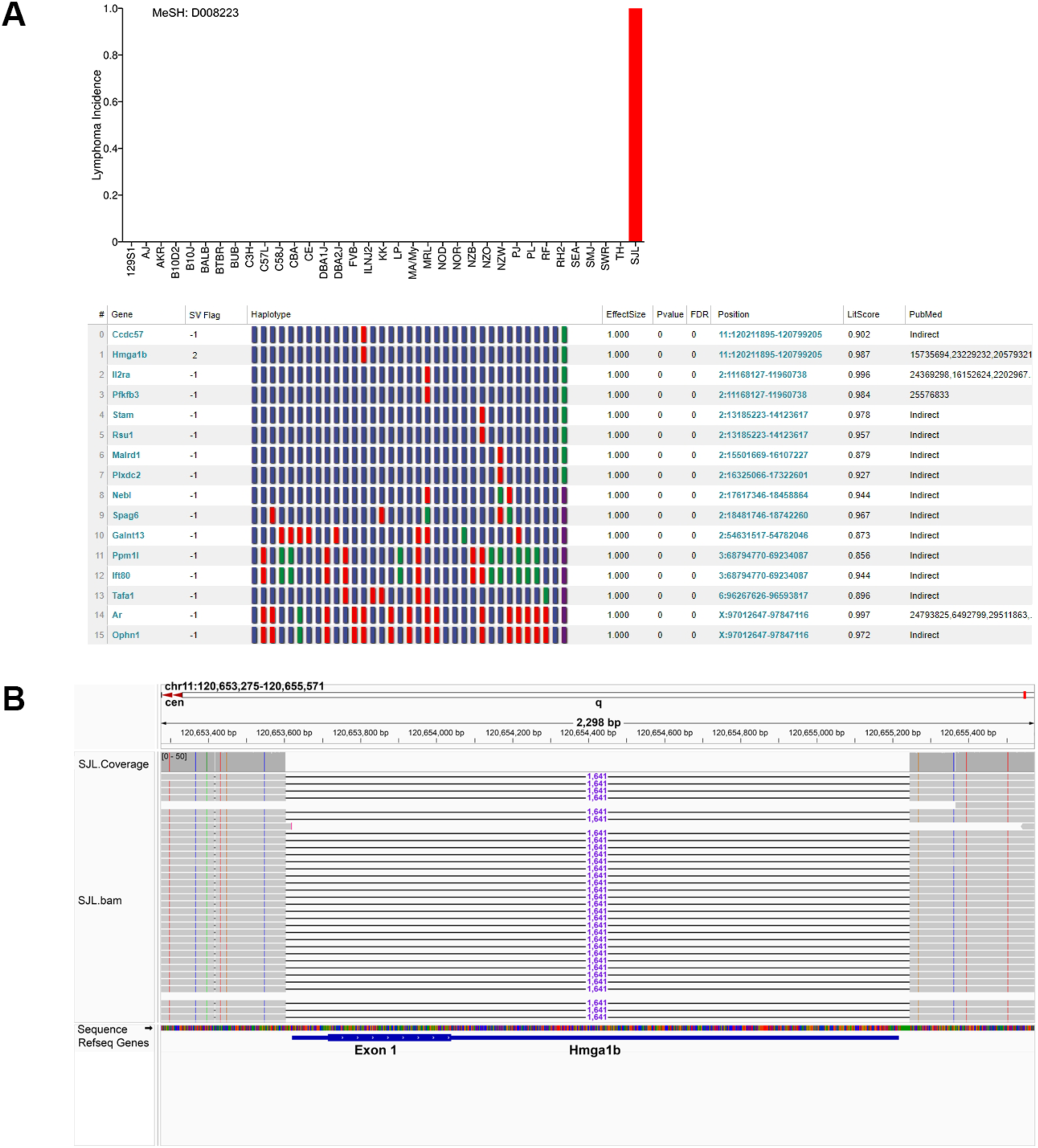
The AI pipeline identifies a genetic factor for lymphoma susceptibility in SJL mice. (A) *Top panel:* Lymphoma was treated as a qualitative trait, which appears in SJL mice (incidence = 1), and not in the 34 other classical inbred strains (incidence = 0). *Bottom panel:* The AI pipeline performed a GWAS to identify haplotype blocks with allelic patterns that corresponded with lymphoma susceptibility in the 35 strains, and then selected those with haplotype blocks that contained a large deletion SV that was only present in SJL ((i.e., genetic effect size = 1 and genetic association p-value = 0). The 16 genes (indicated by gene symbol) meeting these criteria are shown. Within the haplotype box: each block color represents a haplotype for one strain, strains with the same haplotype have the same color, and the blocks are shown in the same strain order as in panel A. The chromosome and the starting and ending position of each haplotype block are also shown. The LitScore represents the strength of the association of each gene with lymphoma as determined by the AI-mediated literature search (MeSH term: Lymphoma D008223). PubMed identification numbers are only provided for genes that have a direct link with the MeSH term. Otherwise, the AI indicates that the gene has an indirect association, which results from MeSH term relationships identified with other proteins that are associated with the gene candidate. The SV flag indicates the impact of the SV deletion as determined by VEP analysis: high impact, 2; and modifier -1. Only *Hmga1b* has a high impact deletion and is directly associated with lymphoma. (B) Among the 35 classical inbred strains, SJL mice uniquely have a 1641 bp deletion within *Hmga1b*. This large deletion (as visualized using the integrative genomics viewer) removes the *Hmga1b* exon that encodes the entire 107 amino acid protein.

Two murine genes encode nearly identical HMGA1 proteins ^48^. *Hmga1b* on chromosome 11 encodes a 107 amino acid protein. *Hmga1* on chromosome 17 generates two predominant mRNAs that produce: a 96 amino acid protein (whose sequence is identical to Hmga1b except 11 amino acids are deleted; or a 107 amino acid protein whose sequence is identical to Hgma1b, which arises by differential splicing (**Figure 6A**). Analysis of *Hmga1* or *Hmga1b* mRNAs in SJL liver and spleen tissue by RT-PCR amplification indicated that the SJL mRNAs are identical to those present in other strains. In contrast, the level of expression of *Hmga1 (or Hmga1b)*-derived mRNAs in SJL thymus are greatly reduced relative to those in the thymus of other strains (**Figure 6B**). Because the 3’ UTRs of *Hmga1-* and *Hmga1b-*encoded mRNAs have a sequence difference at a corresponding site, transcript sequencing was used to identify the gene(s) that encoded these mRNAs in different C57BL/6 tissues. The mRNAs in spleen, liver, bone marrow, kidney and lymph nodes were *Hmga1* encoded, while the thymic mRNAs were *Hmga1b* encoded (**Figures 6C and S8**). Hence, *Hmga1b* mRNA expression predominates in a tissue where lymphocyte development occurs. HMGA protein family members are strongly associated with leukemia and lymphoma in mice ^49^ and humans ^47^, which explains why reduced HGMA protein function in the SJL thymus contributes to lymphoma susceptibility (discussed below).

**Figure 6.**
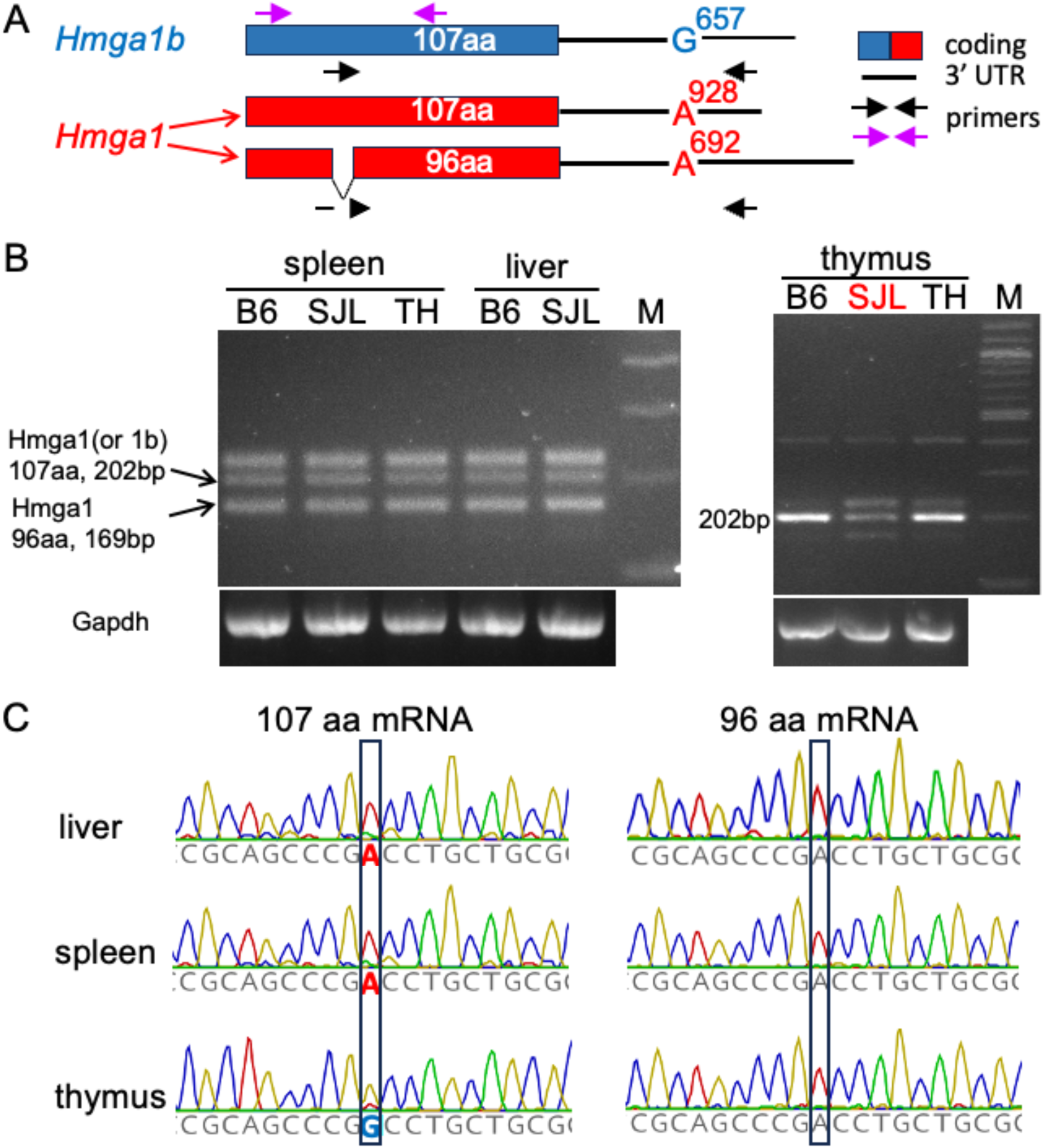
Hmga1b is selectively expressed in the thymus. (A) A diagram of the *Hmga1* and *Hmga1b* encoded mRNAs. *Hmga1b* encodes a 1626 bp mRNA, which generates a 107 amino acid protein. *Hmga1* generates two principal mRNAs: a 1045 bp mRNA that produces a 107 amino acid protein whose sequence is identical to that of Hmga1b; and a 1631 bp mRNA that produces a 96 amino acid protein whose sequence is identical to Hmga1b, except for the deletion of a 11 amino acid segment. The site of a sequence difference at corresponding sites in the 3’ UTR of the *Hmga1* (A) *Hmga1b* (G) mRNAs is shown. The purple arrows indicate the primers used for PCR amplification of the bands shown in B, and black arrows are the primers used to generate the amplicons used for sequencing in C. (B) RT-PCR was performed on spleen, liver and thymic tissues obtained from C57BL/6 (B6), TallyHo (TH) and SJL mice. The amplicons from the *Hmga1* (96 amino acid protein) and the 107 amino acid *Hmga1* (or *Hmga1b*)-encoded proteins are indicated by arrows. The pattern of amplicon expression in liver and spleen is similar in all three strains. In contrast, thymic tissue primarily expresses the 107 amino acid protein, SJL thymus has a much lower level of expression this protein. (C) The thymus only expresses *Hmga1b* mRNA. mRNA was prepared from spleen, liver and thymic tissue obtained from C57BL/6 mice. RT-PCR amplicons from *Hmga1* and *Hmga1b* mRNAs were prepared and sequenced. The 3’ UTR of *Hmga1* mRNA has an A, while *Hmga1b* mRNA has a G at the boxed position. The results indicate that spleen and liver mRNAs are encoded by *Hmga1*, while thymic mRNAs are encoded by *Hmga1b*.

## DISCUSSION

This dataset represents the most comprehensive and carefully performed analysis of SVs in the mouse genome. High quality LRS with a high level of genome coverage for 40 inbred strains, and recently developed state of the art programs (Sawfish ^18^, Sniffles2 ^16^, Cue ^20^) that were adapted for LRS analysis were used with earlier programs (PBSV, Dipcall) for SV identification. These programs were selected based upon simulation results that indicated that they performed optimally for identification of a certain type/size of SV. Large INS and all DUPs were validated by a separate program, and all INVs were individually validated by manual observation. Our results are consistent with prior observations that no single SV calling algorithm was optimal for detection of the different sizes and types of SVs, and that there can be a high level of divergence when the results of simulated and actual data are compared ^39^. Our data also demonstrates that there is a critical need for improved methods for analysis of DUPs and INVs, which probably will require machine learning based programs. Nevertheless, since murine DUPs and INVs were particularly hard to identify, even when LRS was used ^14^; the INV and DUP, along with the INS and DELs identified here could facilitate many genetic discoveries. DUPs and INVs are already known to contribute to Autism ^50,51^ and to impact brain function ^50^. Recent analyses have indicated that segmental duplications occupy ∼7% of the human genome ^52^. One study limitation is that the currently used SV callers were designed and evaluated using human genomic sequences. Since the genomic sequences of the inbred strains differ from outbred human populations, the existing programs may need to be further refined for mouse genomes. Another study limitation is that tandem repeats (**TRs**) – a type of SV with multiple repeats of short DNA sequence motifs, which are associated with multiple human diseases ^53,54^ - were not covered here; but will be analyzed in a subsequent paper. Also, the increased variability between the genomic sequences of the wild-derived and classical inbred stains could also limit the ability of alignment-based programs for SV identification.

The discovery of the second genetic susceptibility factor for SJL lymphoma was facilitated by this murine SV database and the AI genetic discovery pipeline. A novel bi-genic model explains why B cell lymphomas uniquely develop in SJL mice, and this model is consistent with all available data about its pathogenesis. The first genetic factor is a protein (vSAg29) encoded by an endogenous retrovirus in SJL mice ^26,27^ that stimulates CD4 T cells to produce cytokines ^28^ required for B cell lymphoma development ^29^. The second genetic factor is an SJL-unique SV generates an *Hmga1b* KO. This SJL-unique genetic factor was predicted by murine intercross experiments, which indicated that a genetic factor present in multiple other strains suppressed lymphoma development ^31,32^. The *Hmga1b* deletion SV allele is also present in 3 wild derived strains (CAST/Ei, SPRET, MOLF); and the endogenous retrovirus is present in another inbred strain (MA/My); but these strains do not spontaneously develop lymphomas. Hence, a unique combination of two genetic factors (vsAg29 and the *Hmga1b* deletion SV) is required to produce lymphomas in SJL mice. Hopefully, the genetic architecture of other murine genetic models of important biomedical traits will be uncovered using this SV database.

### How could a SJL SV allele that ablates Hmga1b promote lymphoma development?

While Hmga1 and Hmga1b have virtually identical protein sequences, our data demonstrates that only *Hmga1b* mRNAs are expressed in the thymus. A series of *in vitro* studies demonstrated that Hmga1 represses the expression of the Recombination activating gene 2 protein (RAG2) endonuclease ^55^, which plays a key role in lymphocyte development; but increased RAG expression can cause DNA damage and an increased risk for oncologic transformation ^56^. *Hmga1* knockout (KO) mice have increased RAG2 activity in their spleens^55^. An *Hmga1* KO altered T cell development, increased B cell development, and caused the mice to develop B cell lymphomas and other hematopoietic malignancies ^57,58^. SJL spleen and lymph node tissues have an increased number of germinal centers with IL-21 producing T follicular helper (TfH) cells, an increased level of IL-21 production; and SJL lymphoma development is IL-21 dependent ^29^. Hence, the HMGA tumor suppressor function is (at least partly) mediated through repression of RAG2; loss of this repressor function in the SJL thymus will alter T cell development and could induce DNA translocations in other tissues. The abnormalities in thymic T cell development along with VsAg29-induced T cell proliferation explains why SJL (and *Hmga1* KO) mice have an increase in IL-21 producing TfH cells in their germinal centers, which leads to the development of a population of B cells with a high level of abnormal IgH rearrangements. By this mechanism, the activity of an expanded and abnormal population of TfH cells generates an IL-21 dependent B cell lymphoma in SJL mice ^55,58^.

Understanding the genetic architecture of SJL lymphoma susceptibility could provide new insight into the pathogenesis of certain types of human lymphomas. For example, SJL lymphomas have transcriptomic and phenotypic similarities with one type of non-Hodgkin human lymphoma: angioimmunoblastic T-cell lymphoma (AITL) ^24,29^. Like SJL lymphomas, AITL develops late in life, is driven by IL-21-producing TfH cells ^59^, and it sometimes develops into a B cell lymphoma ^60^. Analogous to the role of vSAg29 and the *Hmga1b* deletion allele in driving lymphomas in SJL mice, Epstein Barr virus (EBV) is detected in 66-86% of AITL patients ^61,62^ and mutations in genes encoding epigenetic modifiers have frequently been detected in AITL patients ^24^. Also, IL-21 induces the expression of EBV latent membrane protein 1 (LMP1) in human B cells ^63^, which has been shown to provide survival signals for B cells ^64^ and can rescue transformed cells from apoptosis ^65^. Given the similarities in their pathogenesis, additional studies on SJL lymphomas could provide new information about how infectious agents and host genetic factors jointly contribute to the pathogenesis of AITL, and possibly other types of lymphomas.

#### Limitations of the study

Despite the use of deep learning and alignment-based approaches to identify inversions and duplications, the low concordance among SV identified by the different methods highlights the need to develop more sophisticated algorithms for identifying these types of SV. We used VaPoR for validation of duplications, but many duplications occurring in structurally complex regions failed to validate, which provides additional evidence of the need to develop high-throughput tools for confirmation of inversions and duplications. Finally, reliance on a single reference genome introduces alignment and reference bias, which makes it more difficult to identify novel insertions or strain-specific sequence differences.

Collectively, these limitations highlight the importance of using high fidelity long read sequencing platforms and multiple programs for SV-detection and validation; and we need to develop reference sequences assemblies for multiple strains to capture the full spectrum of structural variation among inbred mouse strains.

## RESOURCE AVAILABILITY

### Lead contact

Requests for further information and resources should be directed to and will be fulfilled by the lead contact, Gary Peltz.

### Materials availability

This study did not generate new unique reagents.

### Data and code availability

- The SNP, INDEL and SV data have been deposited at the Mouse Phenome Database GenomeMUSter at https://mpd.jax.org/genotypes.
- The long-read sequencing (LRS) data have been deposited at the NCBI BioProject database under accession number PRJNA1250604 and are publicly available as of the date of publication.
- All software and analytical methods used in this study are publicly available, as listed in the key resources table.
- Any additional information required to reanalyze the data reported in this paper is available from the lead contact upon request.

## ACKNOWLEDGMENTS

This work was supported by NIH awards (1R01DC021133 and 1 R24 OD035408) to G.P.; and NHGRI Award R01HG012467 to V.P. The funder had no role in the writing of this paper. We thank Dr. Laura Reinholdt (Jackson Labs) for supplying DNA obtained from several of the inbred strains.

## AUTHOR CONTRIBUTIONS

Conceptualization, G.P.; methodology, W.R., F.Q., V.P. and G.P.; formal analysis, software, and validation, W.R., E.D., C.T.S., Z.C., V.P.; visualization, W.R.; writing—original draft, W.R. and G.P.; writing—review & editing, W.R. and G.P.; funding acquisition, V.P. and G.P.; supervision, G.P.

## SUPPLEMENTAL INFORMATION

**Document S1. Figures S1-S8 and Tables S1,S2,S3,S5,S6 and S12**

**Table S4.** Lists of SVs identified in our study and those identified in a prior study. The data are organized into four categories: deletions (DEL), insertions (INS), inversions (INV), and duplications (DUP). The lists include common variants shared between the studies and unique variants that are specific to each dataset.

**Table S7.** Lists of high impact DELs present in all 39 or in the 35 classical inbred strains. The chromosome, starting and ending position, size, gene symbol, strains with the variant allele, and the predicted effect for each DEL are shown.

**Table S8.** Lists of high impact INS present in all 39 or in the 35 classical inbred strains. The chromosome, starting and ending position, size, gene symbol, strains with the variant allele, and the predicted effect for each INS are shown.

**Table S9.** Lists of INVs present in all 39 or in the 35 classical inbred strains. The chromosome, starting and ending position, size, gene symbol, strains with the variant allele, and the predicted effect for each INV are shown.

**Table S10.** Lists of high impact DUPs present in all 39 or in the 35 classical inbred strains. The chromosome, starting and ending position, size, gene symbol, strains with the variant allele, and the predicted effect for each DUP are shown.

**Table S11.** List of DELs present in all 39 or in the 35 classical inbred strains with CTCF binding sites. The chromosome, starting and ending position of the CTCF recognition element and its sequence, size, gene symbol, strains with the variant allele for each DEL, the CTCF score (the strength of the motif match based on the position weight matrix), p-value and Q-value (p-value that is adjusted for the false discovery rate) are shown. These were determined using the CTCF (v0.99.11) R package in Bioconductor. All predicted sites have a *P*-value that is below the recommended cutoff (1 x 10^-6^). The number of peaks in the 42 ENCODE files with CTCF ChIP-Seq data obtained using mouse cells or tissues, which indicates that CTCF is bound to that site, and the data source for each peak is shown.

## STAR★METHODS

### KEY RESOURCES TABLE

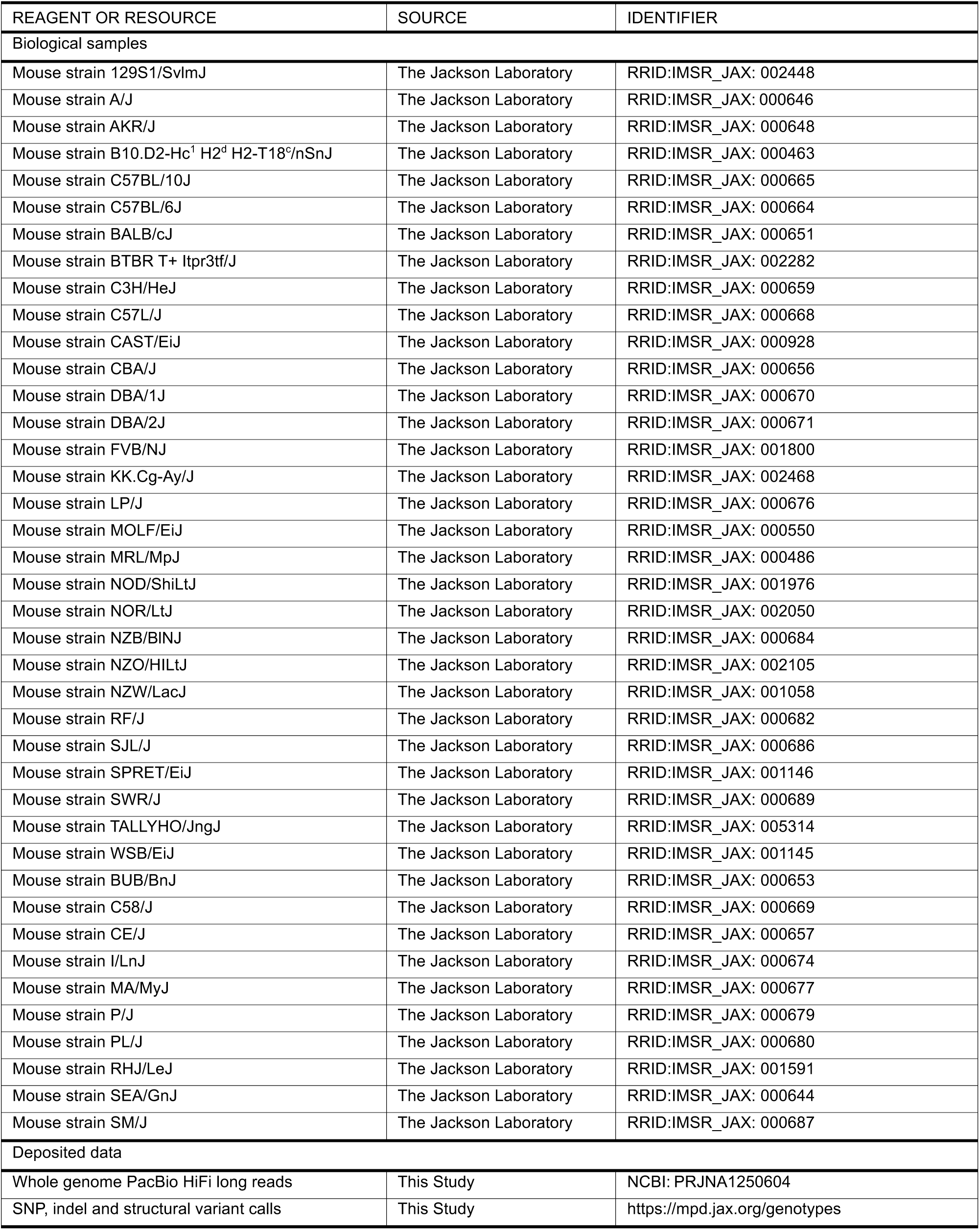
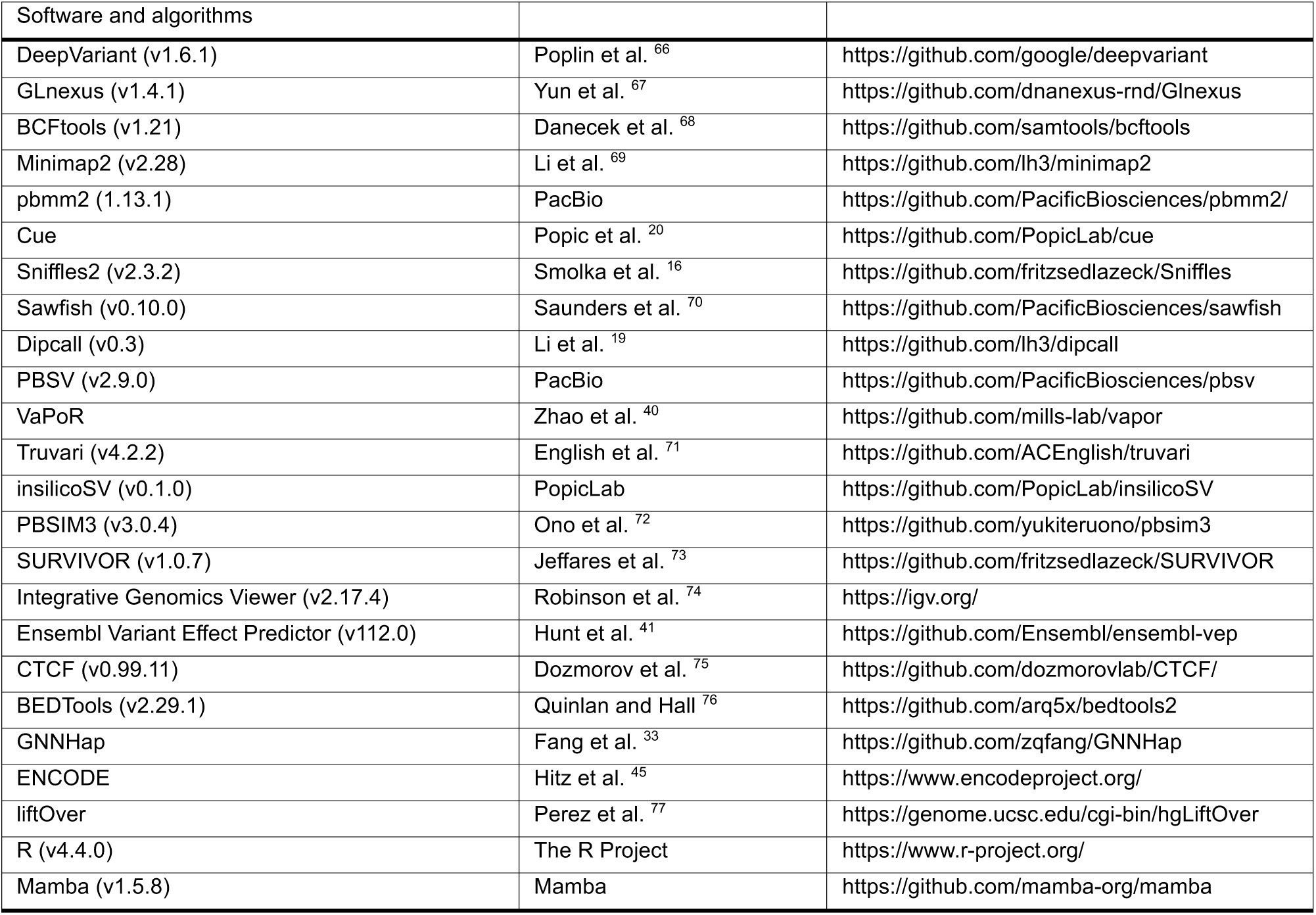

### METHOD DETAILS

#### Animal experiments

All animal experiments were performed according to protocols that were approved by the Stanford Institutional Animal Care and Use Committee. All mice were obtained from Jackson Labs, and the results are reported according to the ARRIVE guidelines ^78^.

#### DNA sequencing

Forty inbred strains (Table S1) were subject to LRS using the HiFi REVIO system (PacBIO). For thirty strains, mouse liver was obtained from mice purchased from the Jackson Laboratory, snap-frozen in liquid nitrogen and shipped on dry-ice to the DNA Technologies Core of the Genome Center, University of California Davis were high molecular DNA purification and REVIO sequencing was performed. The genomic DNA for ten strains (the bottom 10 listed in Table S1) were kindly provided Dr. Laura Reinholdt, Co-Director of the Mutant Mouse Resource and Research Center at the Jackson Laboratory (Bar Harbor, ME); and REVIO sequencing was also performed at the UC Davis Genome Center.

#### Sequencing Data Quality Control

Our study utilized PacBio HiFi sequencing technology, which is known for its high accuracy in single- molecule real-time (SMRT) sequencing. The quality control metrics for our sequencing data include:

1. Accuracy Yield: We achieved high-quality bases (>QV20/99% accuracy) with a yield typically ranging between 75-105 Gb, averaging around 90 Gb on the Revio system.
2. Fragment Size: The SMRTbell library was prepared with an optimal fragment size of 15-20 kb for HiFi whole-genome sequencing.
3. Read Length: The mean HiFi read length was between 15-20 kb.
4. Read Quality: The median HiFi read quality was Q30, with a range from Q28 to Q32.
5. Sequencing Control: We employed a 110 kb control sequence, with typical control read lengths around 70 kb (ranging from 50-100 kb) and counts exceeding 1000.

#### Identification of SNPs and INDELs

PacBio HIFI long read sequence (LRS) data were aligned to GRCm39 (mm39) reference genome using minimap2 (v2.28) ^69^ and pbmm2 (v1.13.1) to generate the BAM files. SNP and INDEL alleles were identified for each strain using the GPU-based DeepVariant (v1.6.1) ^66^ and the alleles from all strains were merged using GLnexus (v1.4.1) ^67^.

#### Alignment, Variant Filtering and Quality Control for SNPs and INDELs

After alignment, SNPs and INDELs were identified using DeepVariant, they were consolidated with GLnexus. Then, a series of stringent filtering criteria were applied using BCFtools (v1.21) ^68^ to ensure the reliability of our variant calls.

1. Sequence Mapping: We employed two aligners—minimap2 and pbmm2—to map the sequences. For example, for the 129S1 strain, the following commands were used:

For minimap2:

minimap2 -c -a --MD -x map-hifi -t 96 /path1/mm39.fa /path2/129S1.fastq.gz > /path3/129S1.align.bam For pbmm2:

pbmm2 align /path1/mm39.fa /path2/129S1.fastq.gz /path3/129S1.align.sort.pbmm2.bam --preset HIFI -- sort --rg ‘@RG\tID:129S1\tSM:129S1’ -j 64

These parameters ensure high-quality alignment by applying options that are optimal for PacBio HiFi reads.

2. Variant Calling for SNPs/INDELs. We used the GPU version of DeepVariant with default options tailored for PacBio data. For instance, the DeepVariant command was executed as follows: docker run --name 129S1 --gpus ‘“device=0”’ -v “/path1/input”:“/input” -v “/path2/output”:“/output”

google/deepvariant:“1.6.1-gpu” /run_deepvariant --model_type=PACBIO --ref=/input/path3/mm39.fa -- reads=/input/minimap2/129S1.align.sort.bam --output_vcf=/output/129S1.raw.mini.vcf.gz -- output_gvcf=/output/129S1.g.raw.mini.vcf.gz --intermediate_results_dir

/output/129S1_intermediate_results_dir --num_shards=32 --sample_name=129S1 And then:

glnexus_cli --config DeepVariant *.g.raw.mini.vcf.gz > deepvariant.bcf --dir scratch --trim-uncalled-alleles -

-threads 96

3. Chromosomal Regions: Filtered to include only autosomal chromosomes (chr1 to chr19) and the X chromosome.

4. Filter Status: Included variants where the FILTER field is marked as “PASS”.

5. Quality Score: Excluded variants with a QUAL score below 30.

6. Depth and Genotype Quality: Retained variants with a read depth (DP) of at least 10 and a genotype quality (GQ) of at least 30.

7. Genotype Call: Selected variants with a homozygous alternate genotype (“1/1”).

8. Variant Type: Filtered to retain only biallelic SNPs and INDELs.

#### Simulations for assessing SV program performance

We generated 8 synthetic genomes using insilicoSV (https://github.com/PopicLab/insilicoSV), which inserted 1000 simulated SVs of each type and size into the GRCm39 reference sequence. For each synthetic genome, PacBio HiFi reads at 30x coverage using PBSIM3 (v3.0.4) ^72^ were simulated, and the simulated reads were aligned using minimap2.

#### Assessing SV identification programs

Since SV analysis programs vary in their ability to identify different types of SVs ^38,39^, we performed a set of targeted simulations to examine the ability of five programs (Cue ^20^, Sawfish (v0.10.0) ^70^, Sniffles2 (v2.3.2) ^16^, PBSV (v2.9.0), and Dipcall (v0.3) ^19^) to detect SVs that were artificially inserted into the mouse genomic sequence. We separately assessed their ability to identify small (<1 kb) and large (1-100 kb) DELs, INSs, INVs and DUPs (Table 1). This strategy measures the upper bound on recall for each program and SV category to identify those that systematically miss specific types of SVs. Multiple methods had high recall (>90%) for small and large DELs (Cue, Sniffles2, Sawfish and PBSV), large DUPs (Cue, PBSV), large INVs (Cue, Sniffles2 and PBSV), and small and large INS (Cue, PBSV).

However, only Dipcall (which used individually assembled genomic sequences) could reliably detect large INS.

The simulation results were used to select the SV identification programs and database curation methods used to assemble the SV database (Figure 1). A recent analysis of human LRS data revealed that SVs found by only one program are more likely to be false positives ^38^. Therefore, a consensus-based approach was used for SV types where multiple programs exhibited high recall in the simulations, and where a high level of agreement was observed when the real data was analyzed by these programs.

Also, whenever possible, we used a machine learning-based method (Cue) with a heuristic-based method (Cue, Sniffles2, PBSV), since programs that use similar heuristics may jointly produce false positives. For example, 85% of the small and 74% of DELs were commonly identified by Cue, Sawfish and Sniffles2, which indicates that these SVs were valid (Figures 2A and S5A). However, if only a single caller achieved high recall in the simulations (i.e., large INSs), or when low agreement was observed among the programs when the actual data was analyzed (i.e., INVs) (Figures 2A and S5A), the consensus strategy was replaced with one where the individual predictions obtained from one or more high recall tools were selected if they could be validated by a computational program or visual inspection. We used VaPoR ^40^, which is a program that autonomously validates SVs identified by analysis of LRS data, to identify the high-quality SVs. For example, many of the validated INVs were only identified by two of the three analysis programs utilized, and most small and large DUPs were only identified by one program (Figures 2A and S5A). The low level of overlap supports our strategy of using different methods but only incorporating SVs, which were validated by another method, into the database. We are aware that the VaPoR validation requirement is likely to eliminate some true positive SVs, but we accept this risk to ensure that only true positive SVs were included in the database.

The simulation results for Sniffles2 and Sawfish for inversions and small DUPs were withheld due to performance issues that became apparent while evaluating the results produced by those programs. First, we examined why the simulations for recognition of small DUPs by Sniffles2 and Sawfish produced low values for precision, recall and F1 (W^$^). By changing the parameters [typeignore=true] used by the Truvari (v4.2.2) ^71^ software to calculate the precision, recall and F1 values, we found that small DUPs were incorrectly labelled by these programs as INS. Although we used IGV images to visually validate the high impact INS, it is possible that some DUPs were labelled as INS. Second, we also found a bug that was specifically present in the early version of Sawfish that was used in this study interfered with the recognition of INVs (W*).

#### Identification of SVs

To identify small and large deletions, Cue and Sniffles2 were used to perform SV calling based on the minimap2 alignment, and Sawfish was used for SV calling based on the pbmm2 alignment. Then, SURVIVOR (v1.0.7) ^73^ was used to merge the results obtained by these three programs. The SVs commonly identified by at least two of these programs were included in the final dataset. Sniffles2 and Sawfish were used to identify small insertions; the small insertions jointly identify by both programs were merged using SURVIVOR and were incorporated into the final dataset. An assembly-based program Dipcall was used to identify large insertions, and VaPoR was used for their validation. The large INS with a VaPoR statistic GS≥0.15 and QS ≥ 0.1 were considered to be validated, and were incorporated into the final dataset. Cue, Sniffles2 and PBSV were used to identify INVs: the Cue and Sniffles2 analyses were based on minimap2 alignments, whereas the PBSV analysis was based on pbmm2 alignment. SURVIVOR was used to merge those identified by any of the three methods; these INVs were then inspected using the Integrative Genomics Viewer (IGV) (v2.17.4) ^74^ as described below; and visually- validated INVs were incorporated into the dataset. DUPs were identified by PBSV or Cue, and then validated by VaPoR as described above. In addition, the starting and ending positions of the identified DUPs were examined to determine if they were located within an identified insertion. Functional annotation was performed using the Ensembl Variant Effect Predictor (VEP) (v112.0) ^41^.

#### SV Variant Filtering and Quality Control

Additional information about sequence quality, read support, genotype calls and quality control metrics used for SV identification.

1. Sequence Mapping Quality. For Cue, Sniffles2, Sawfish, and Dipcall, we applied a filter of QUAL > 30 to ensure high mapping quality. Note that PBSV 7 does not provide a corresponding parameter for this metric.
2. Read Support for Each Variant. For Sniffles2, we required a minimum read support of SUPPORT ≥ 10 (where SUPPORT represents the number of reads supporting the SV).

For PBSV, we applied a read depth filter of DP ≥ 10 (with DP indicating the read depth at the variant position)

3. Genotype Quality. Both Sawfish and Sniffles2 required a genotype quality (GQ) of at least 30 to retain a variant call.

4. Additional Filtering using BCFtools for all methods. We further refined the SV calls by applying the following criteria:

(1). Variants were restricted to the 19 autosomes and the X chromosome
(2). Only variants with FILTER = “PASS” were retained.
(3). Variants with an SV length (SVLEN) between 50 and 500,000 bp were included.
(4). Variants annotated as SVTYPE = “BND” were excluded.
(5). Genotype calls of “0/0” were removed.

#### Visual inspection using IGV

Visual inspection of high impact deletions and insertions, and all inversions was performed by examining IGV images. To do this, the IGV image script code was modified as follows:

“new
genome mm39
snapshotDirectory /pathway1/
load /pathway2/name.identify.vcf
load /pathway3/name.alignment.bam
goto chr*:start-end position
scrollToTop
preference SAM.COLOR_BY READ_STRAND
preference SAM.LINK_READS TRUE
preference SAM.LINK_TAG READNAME
preference SAM.GROUP_OPTION ZMW
preference SAM.SHOW_MISMATCHES TRUE
preference SAM.MAX_VISIBLE_RANGE 500
snapshot output.png
exit”

We found that altering three of the six display settings was important. (i) The ‘preference SAM.COLOR_BY READ_STRAND’ indicates that reads from different strands are displayed in different colors. (ii) The ‘preference SAM.GROUP_OPTION ZMW’ specifies the ZMW group option, which is optimal for analysis of PacBio long read sequence. (iii) The ‘preference SAM.MAX_VISIBLE_RANGE 500’ sets the maximum length of a displayed SV at 500 KB. The use of the alternate parameters improved the display of the inversions. Of note, we used split-read alignments with strand flips when validating INVs by visual inspection, but this approach has limitations since it can only validate INVs where split-read alignments are available.

#### Predicting CTCF binding sites

The predicted CTCF binding sites in the genomic sequences of the inbred strains were identified using the CTCF (v0.99.11) ^75^ R package in Bioconductor with the JASPAR 2022 database and the recommended *P*-value cutoff (1 x 10^-6^). We then examined the functional relevance of these predicted CTCF binding sites using ChIP-Seq data for CTCF from the ENCODE project ^45^. Specifically, we obtained 42 narrowPeak BED files with ChIP-Seq experiments conducted on various mouse tissues or cell types, which had the assay type designated as ‘DNA binding,’ the assay title as ‘TF ChIP-seq,’ and the assay target as ‘CTCF.’ These datasets were aligned to the GRCm38 reference genome. To ensure consistency, we first converted our predicted CTCF binding site coordinates from GRCm39 to GRCm38 using the Lift Genome Annotations tool ^77^. For the ChIP-Seq data, we applied stringent filtering criteria, selecting peaks with a signal intensity of at least 50 and a -log10(q-value) greater than 3 to ensure that high quality data was analyzed. We then utilized the BEDTools (v2.29.1) ^76^ suite to compare our predicted CTCF binding sites with the ChIP-Seq peaks. A CTCF site was considered as bound if at least 50% of its length overlapped with a ChIP-Seq peak. We then recorded the number of supporting peaks obtained from the 42 datasets and documented their respective sources.

#### SNP and SV dataset comparisons

To compare the SNPs identified here using the PacBio HiFi long-read sequencing (LRS) with the previously identified SNPs, which were identified using short-read sequencing (SRS) ^5^, we selected 39 strains that were found in both datasets. The SRS variants calls underwent the same variant filtering process as the LRS SVs. Since the SRS results were aligned to the GRCm38 reference genome, the liftOver tool was used to convert their genomic coordinates to the GRCm39 reference. For comparison, we generated unique variant identifiers by concatenating the chromosome, position, reference allele, and alternative allele for each SNP. These variant IDs were then compared using the ‘intersect()’ and ‘setdiff()’ functions in R, which enabled us to identify the shared and unique variants in the two datasets.

The SVs identified in our 39 inbred strain panel were compared with those identified by the Jackson Laboratory’s analysis of 14 inbred strains ^15^. For this analysis, deletions (DEL), insertions (INS), duplications (DUP), and inversions (INV) were separately analyzed. Comparisons were performed using a position tolerance-based approach. SVs from different datasets were considered to represent the same variant if their starting and ending positions, and their lengths were less than 50 bp. If any of these conditions were not met, the SVs were considered as distinct. The IDs of the shared and unique SVs in the two datasets were reported with each ID formatted as ‘chromosome-start_position-SV_type- SV_length’ (e.g., ‘chr1-3101582-DEL-357’) in Table S4.

#### Analysis of Hmga1b and Hmga1 mRNAs

Total RNA was purified from mouse thymus, spleen and liver obtained from SJL, C57BL/6 and TallyHo mice; and from kidney and hind leg bone marrow of SJL and C57BL/6 mice using the TRIzol (Thermo Fisher) reagent with the Direct-zol RNA miniprep kit (Zymo Research). One half ug each of the total RNAs were reverse transcribed using the High-Capacity cDNA Reverse Transcription kit (Applied Biosystems) in a 20 μl volume. One μl each of the cDNAs were then subject to PCR using the GoTaq® G2 DNA polymerase master mix (Promega) and primers for *Hmga1b*, *Hmga1* or *Gapdh* (control) transcripts.

*Hmga1b*/*Hmga1* transcript primers: Hmga-F1: AGCGAGTCGGGCTCAAAGTC and Hmga-R1: CGCCCTTATTCTTGCTTCCCTTT. Primers for either the 107aa transcript or the 96aa transcripts were: Hmg-107aa-F1, TGAGTCCTGGGACGGCGCT, Hmg-96aa-F, AGCAGCCTCCGAAAGAGCC, Hmga-R, GAATGCTCCCAGGACCCTCTA. Primers for the *Gapdh* transcript: Gapdh-F2: GTAGACAAAATGGTGAAGGTCGGT and Gapdh-R1, GGTCCAGGGTTTCTTACTCCTTG. The PCR amplified cDNA generated using Hmg-107aa-F1 and Hmga-R (the 107aa transcripts only) primers or the Hmg-96aa-F and Hmga-R (the 96aa transcripts only) primers were gel purified and then subject to Sanger sequencing.

#### Analysis of other genes identified by the mouse genetic AI pipeline

In addition to Hmga1b, only three other genes with SJL-specific large deletions were identified as having a potential direct association with lymphoma. (i) Il2ra was associated with lymphoma because Il2ra mRNA expression is a prognostic marker for Burkitt lymphoma 12, acute myelogenous leukemia 13, mycosis fungoides 14, diffuse large B cell lymphoma 15 and other cancers; and IL-2 is administered as an anti- cancer agent 16. The absence of Il2ra could reduce T and NK cell mediated anti-cancer immunity, which potentially could facilitate B cell lymphoma development. However, the Il2ra SV is not high impact (it labelled as a modifier and does not affect the coding sequence), and will not have a major effect on Il2ra expression. (ii) Although Pfkb2 was identified as having a direct association with lymphoma, the paper 17 identified by the AI was a false positive association with lymphoma. (iii) Ccdc57 identified by the AI because it is within the same haplotype block as Hmga1b; but Ccdc57 has no direct association with lymphoma. Ccdc57 is in the Human Protein Atlas, which contains lymphoma tissue. No other gene with SJL-specific high impact large deletion was directly associated with lymphoma.

